# Tn5Prime, a Tn5 based 5’ Capture Method for Single Cell RNA-seq

**DOI:** 10.1101/217117

**Authors:** Charles Cole, Ashley Byrne, Anna E. Beaudin, E. Camilla Forsberg, Christopher Vollmers

**Affiliations:** Department of Biomolecular Engineering, University of California Santa Cruz, CA; Department of Molecular, Cellular, Developmental Biology, University of California Santa Cruz, CA; The authors wish it to be known that, in their opinion, the first 2 authors should be regarded as joint First Authors; Current Address: Department of Molecular and Cell Biology, School of Natural Sciences, University of California-Merced, Merced, CA, USA; Institute for the Biology of Stem Cells, University of California Santa Cruz, CA

## Abstract

RNA-seq is a powerful technique to investigate and quantify entire transcriptomes. Recent advances in the field have made it possible to explore the transcriptomes of single cells. However, most widely used RNA-seq protocols fail to provide crucial information regarding transcription start sites. Here we present a protocol, Tn5Prime, that takes advantage of the Tn5 transposase based Smartseq2 protocol to create RNA-seq libraries that capture the 5’ end of transcripts. The Tn5Prime method dramatically streamlines the 5’ capture process and is both cost effective and reliable. By applying Tn5Prime to bulk RNA and single cell samples we were able to define transcription start sites as well as quantify transcriptomes at high accuracy and reproducibility. Additionally, similar to 3’ end based high-throughput methods like Drop-Seq and 10X Genomics Chromium, the 5’ capture Tn5Prime method allows the introduction of cellular identifiers during reverse transcription, simplifying the analysis of large numbers of single cells. In contrast to 3’ end based methods, Tn5Prime also enables the assembly of the variable 5’ ends of antibody sequences present in single B-cell data. Therefore, Tn5Prime presents a robust tool for both basic and applied research into the adaptive immune system and beyond.

## Introduction

As the cost of RNA-sequencing has decreased, it has become the gold standard in interrogating complete transcriptomes from bulk samples and single cells. RNA-seq is a powerful tool to determine gene expression profiles and identify transcript features like splice-sites. However, standard approaches lose sequencing coverage towards the very end of transcripts. This reduced coverage means that we cannot confidently define the 5’ ends of mRNA transcripts which contain crucial information on transcription initiation start sites (TSSs) and 5’ untranslated regions (5’UTRs). Analyzing TSSs can help infer the active promoter landscape, which may vary from tissue to tissue and cell to cell. Analyzing 5’UTRs, which may contain regulatory elements and structural variations can help infer mRNA stability, localization, and translational efficiency. Identifying such features can help elucidate our understanding of the molecular mechanisms that regulate gene expression.

The loss of sequencing coverage towards the 5’ end of transcripts is often attributed to how sequencing libraries are constructed. For example, the widely used Smartseq2 RNA-seq protocol, a powerful tool in deciphering the complexity of single cell heterogeneity (1–3), features reduced sequencing coverage towards transcript ends. This lost information is a result of cDNA fragmentation using Tn5 transposase. Several technologies have tried to compensate for the lack of coverage by specifically targeting the 5’ ends of transcripts. The most notable methods include cap analysis of gene expression (CAGE), NanoCAGE, and single-cell tagged reverse transcription sequencing (STRT) (4–7). CAGE uses a 5’ trapping technique to enrich for the 5’-capped regions by reverse transcription (7). This technique is extremely labor intensive and involves large amounts of input RNA. The NanoCAGE and STRT methods target transcripts using random or polyA priming and a template-switch oligo technique to generate cDNA (4, 6). While NanoCAGE can analyze samples as low as a few nanograms of RNA, and STRT can be used to analyze single cells, they both require long and labor-intensive workflows including fragmentation, ligation, or enrichment steps. Therefore, none of the current 5’ end specific protocols are capable of efficiently and cost-effectively processing hundreds to thousands of single cells necessary to understand heterogeneity within complex mixtures of cells present in, for example, the adaptive immune system or cancer.

Furthermore, new droplet based high-throughput single cell RNAseq approaches like Drop-Seq and 10X Genomics Chromium platform can process thousands of cells but can only analyze the 3’end of transcripts due to integrating a sequencing priming site into the oligodT primer used for reverse transcription. By losing information of the 5’ end almost entirely, these approaches are not capable of comprehensively analyzing cells of the adaptive immune cells which express antibody or T cell receptor transcripts featuring unique V(D)J rearrangement sequence information on their 5’ end.

To overcome this lack of high-throughput single cell 5’ capture methods, we chose to modify the Smartseq2 library preparation protocol which is relatively cost-effective and simple with features of STRT which captures 5’ ends effectively. Here we describe a robust and easily implemented method called Tn5Prime that performs genome-wide profiling across the 5’ end of mRNA transcripts in both bulk and single cell samples. The protocol is based on integrating one sequencing priming site into the template switch oligo used for reverse transcription and subsequently tagmenting the resulting amplified cDNA by Tn5 enzyme loaded with an adapter carrying the other sequencing priming site. This combination allows for the construction of directional RNAseq libraries with one read anchored to the 5’ end of transcripts without the need for separate fragmentation, ligation, and, most importantly, enrichment steps. Additionally, by incorporating cellular identifiers into the template switch oligo makes it conducive for pooling samples after reverse transcription, thereby increasing throughput and reducing cost. Finally, data produced by this novel approach allows for the identification of transcription start sites, the quantification of transcripts, and the assembly of antibody heavy and light chain sequences from single B cells at low sequencing depth.

## Results

### Construction of Tn5Prime libraries

Tn5Prime libraries can be constructed from either purified total RNA or single cells sorted by FACS into multiwell PCR plates. Tn5Prime libraries create a directional paired-end Illumina RNAseq library with read 1 anchored to the 5’ end of transcripts. Directionality and read 1 anchoring is achieved through the use of our modified template-switch oligo and custom Tn5 enzyme. After the addition of reverse transcriptase to total RNA or cell lysate, first-strand synthesis occurs using a modified oligo-dT and a template-switch oligo (TSO) containing a partial Nextera A adapter sequence and, optionally, a cellular index sequence (Table S1, Fig. 1A). During reverse transcription, the oligo-dT serves as a primer at the 3’ polyA tail of mRNA transcripts, while the sequence of the partial Nextera A template-switch oligo is attached to the 3’ end of the synthesized cDNA corresponding to the 5’ end of transcript sequences. After reverse transcription, samples with non-overlapping cellular indexes can be pooled. The cDNA product is then amplified using a complete Nextera A primer and a primer complementary to the modified 5’ end of the oligo-dT. After amplification, the cDNA product will contain a complete Nextera A adapter including Illumina indexes. At this point, samples that contain the non-overlapping Illumina indexes can be pooled. By pooling after reverse transcription and PCR amplification, we can dramatically reduce the workflow complexity and reagent usage.

**Fig. 1.**
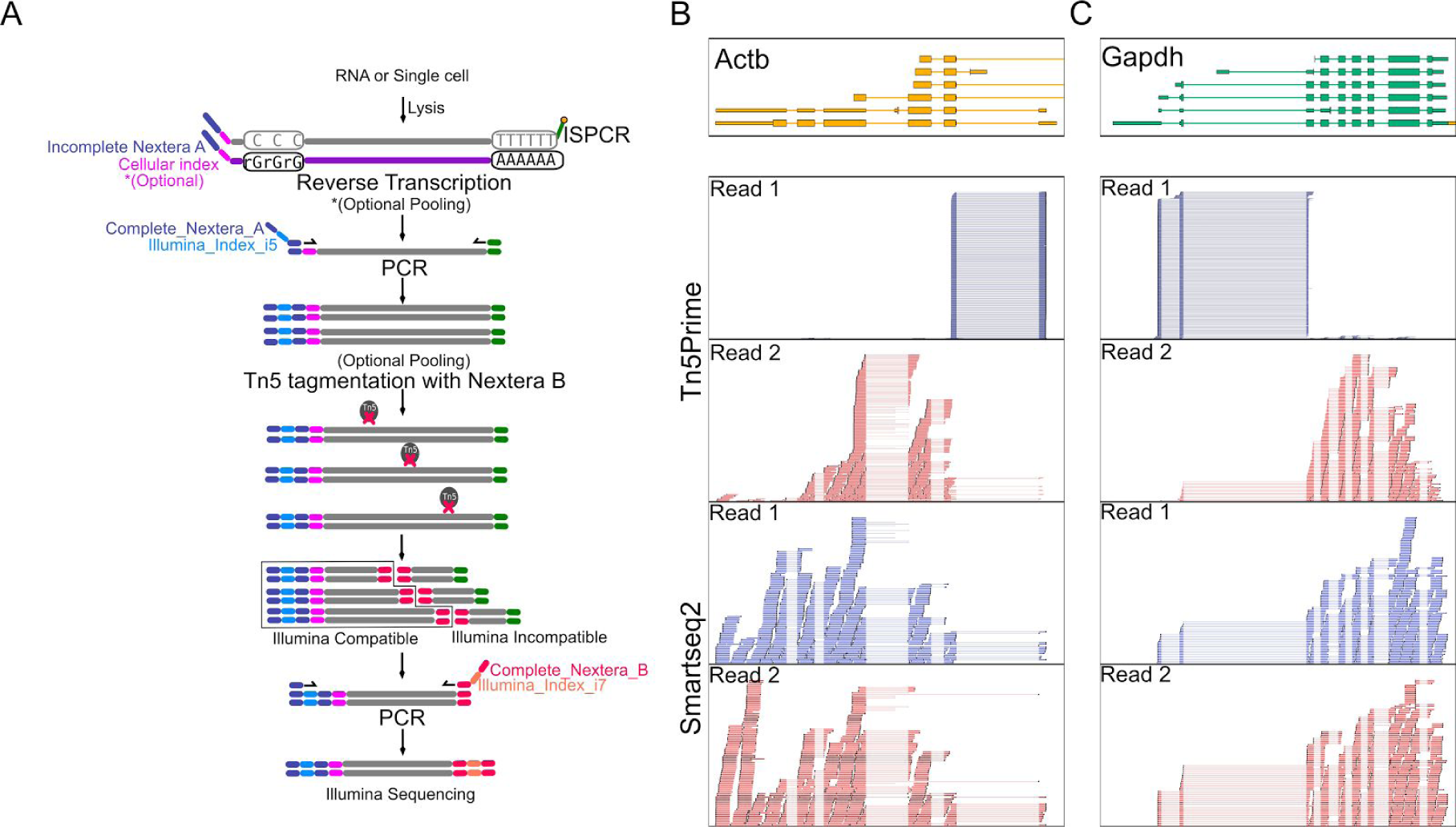
Tn5Prime Library construction and 5’ capture. A.) Schematic of the Tn5Prime library construction. No enrichment steps are required to generate a library that captures the 5’ end of transcripts. B.) Examples of 5’ end capture by Tn5Prime compared to random fragmentation by Smartseq2. Individual alignments for the first (Read1, blue) and second (Read2, red) read of each read pair are shown. Read1 density is shown for both library types as a histogram (blue). Gene models are shown on top (Color indicates transcriptional direction.)

Next, Tn5 transposase, loaded only with a partial Nextera B adapters, fragments the cDNA and attaches the partial Nextera B adapters to the cDNA in a single reaction. The cDNA fragments are then amplified using a universal A primer and a Nextera B primer that primes off the partial Nextera B adapter sequences attached by the Tn5 enzyme. The final product is compatible with the Illumina platform by containing the complete Nextera A and Nextera B adapters. Libraries are then ready to be size selected and quantified prior to sequencing. At this point, no enrichment step is necessary, as only molecules containing both Nextera A and B adapters will be targeted for sequencing. Since only the TSOs associated with the 5’ end of transcripts contain Nextera A adapters, read 1 of all read pairs in the sequencing reaction begins at these 5’ ends and extends into the transcript body, thereby identifying transcription start site and directionality (Fig. 1A-C). Read 2 is distributed throughout the gene body, as each location represents the random insertion of Nextera B adapters by Tn5 and library size selection (Fig. 1B,C).

### Creating and analyzing Tn5Prime data of GM12878 cell line RNA

To evaluate whether our Tn5Prime protocol consistently identifies the 5’ end of the transcript we first performed low coverage RNAseq of total RNA of GM12878 cultured lymphoblast cells. We performed a side-by-side comparison of our protocol with the standard Smartseq2 protocol using the same starting material. Using the HiSeq2500 platform (Illumina) we obtained 570805 and 453761 raw read pairs for two replicate Tn5Prime libraries. We next obtained 1094530 raw read pairs from a Smartseq2 library. Adapter sequences and low quality reads were removed using Trimmomatic (8). In the Tn5Prime replicates, 92.51% and 92.62% of the trimmed and filtered reads mapped uniquely to the human genome using the STAR alignment tool (9), surpassing the standard Smartseq2 protocol at 88.50%. The uniquely aligned reads from the TN5Prime replicates collectively had a redundancy of 1.34. This high unique alignment percentage indicates that our Tn5Prime protocol produces libraries of high complexity.

### Detecting Transcription Start Sites using Tn5Prime

We analyzed the read distribution across transcripts both visually and systematically to determine the 5’ specificity of our protocol. Visual inspection found that while Smartseq2 reads are distributed across the entire body of genes, Tn5Prime reads follow two distinct patterns: First, the start of the read 1 is anchored to the transcription start site. Second, the start of read 2 is variable and likely dependent on size selection during library preparation (Fig. 1B). Next, systematic analysis was based on mapping the start of read 1 to identify putative Transcription Start Sites (TSSs). To test our ability to identify TSSs, we compared our Tn5Prime data to the Gencode genome annotation and CAGE data which was generated from the same GM12878 cell line from the ENCODE project. We identified putative TSSs by calling peaks enriched from the start of read 1 in our Tn5Prime data (see Methods). 89.7% of the 17853 peaks fell within TSSs (0-25 bp upstream) with the vast majority of them falling near promoter regions (26bp-1000bp upstream) or 5’UTRs (Fig. 2A). Next, we subsampled the CAGE data to levels similar to the Tn5Prime data and called peaks in the same manner. 14107 of 17853 Tn5Prime peaks (73%) fell within 25bp to the nearest of 27526 CAGE peaks, indicating high concordance between the two approaches (Fig. 2B). Tn5Prime peaks (3,746) that were not within 25bp of a CAGE peak contained far less sequencing reads on average than those within 25bp of a CAGE peak. These results indicate that these transcripts might be expressed at lower levels and show more variance between the Tn5Prime and CAGE datasets (Fig. 2B). Ultimately, this suggests that our Tn5Prime protocol is equivalent to the gold standard CAGE technique in targeting transcription start sites.

**Fig. 2.**
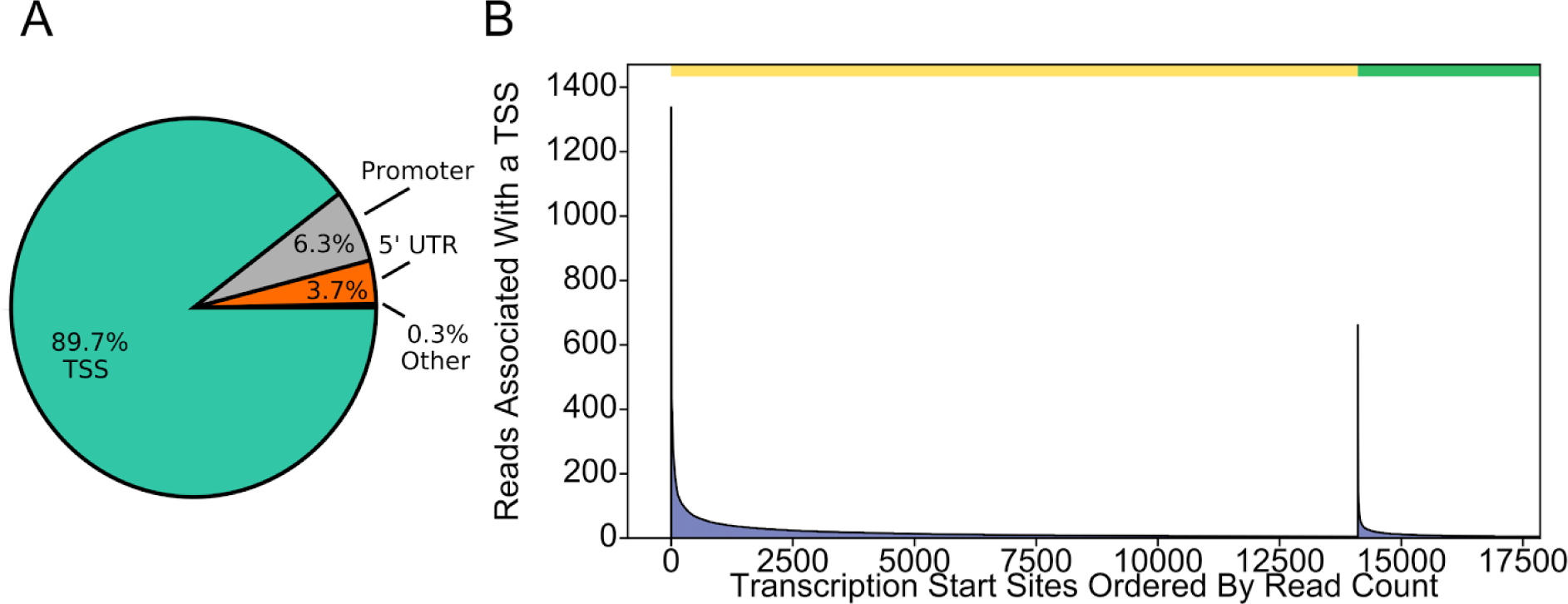
Tn5Prime peaks are highly concordant with GENCODE annotation and CAGE peaks. A) Tn5Prime peaks were matched to features in the Gencode annotation and the feature they matched are shown as a pie chart. B) Tn5Prime were matched to CAGE peaks. The green bar on top indicates the peaks within 25bp and the yellow bar indicates all other peaks. Peaks in each were rank sorted according to their read coverage and shown as a histogram.

### Quantifying the Transcriptome using Tn5Prime

After validating the ability of Tn5Prime to detect transcription start sites, we next wanted to examine whether it is capable of transcript quantification. To determine whether our Tn5Prime method is quantitative we compared GM12878 data generated from four different protocols: Tn5Prime, Smartseq2 data generated by our lab, as well as CAGE and RNA-seq data produced by the ENCODE project (Fig. 3). We used the Tn5Prime data mentioned in the previous section and generated the Smartseq2 data on the same Cell line as described by (1). We performed replicates using the Tn5Prime protocols to define overall reproducibility and accuracy. Based upon our results, transcript quantification by Tn5Prime replicates showed extremely high correlation with a Pearson correlation coefficient of r=0.97 (95% C.I. 0.97-0.97). Quantification by Tn5Prime also correlated very well with Smartseq2 with a Pearson r of 0.87 (95% C.I. 0.86-0.87). Tn5Prime and Smartseq2 data were comparable with ENCODE RNA-seq and CAGE data (Fig. 3), indicating that the Tn5Prime protocol is equivalent to the conventional Smartseq2 method in measuring transcript abundance. Together, these data show that Tn5Prime can accurately identify transcription start sites and quantitatively measure transcript abundance.

**Fig 3.**
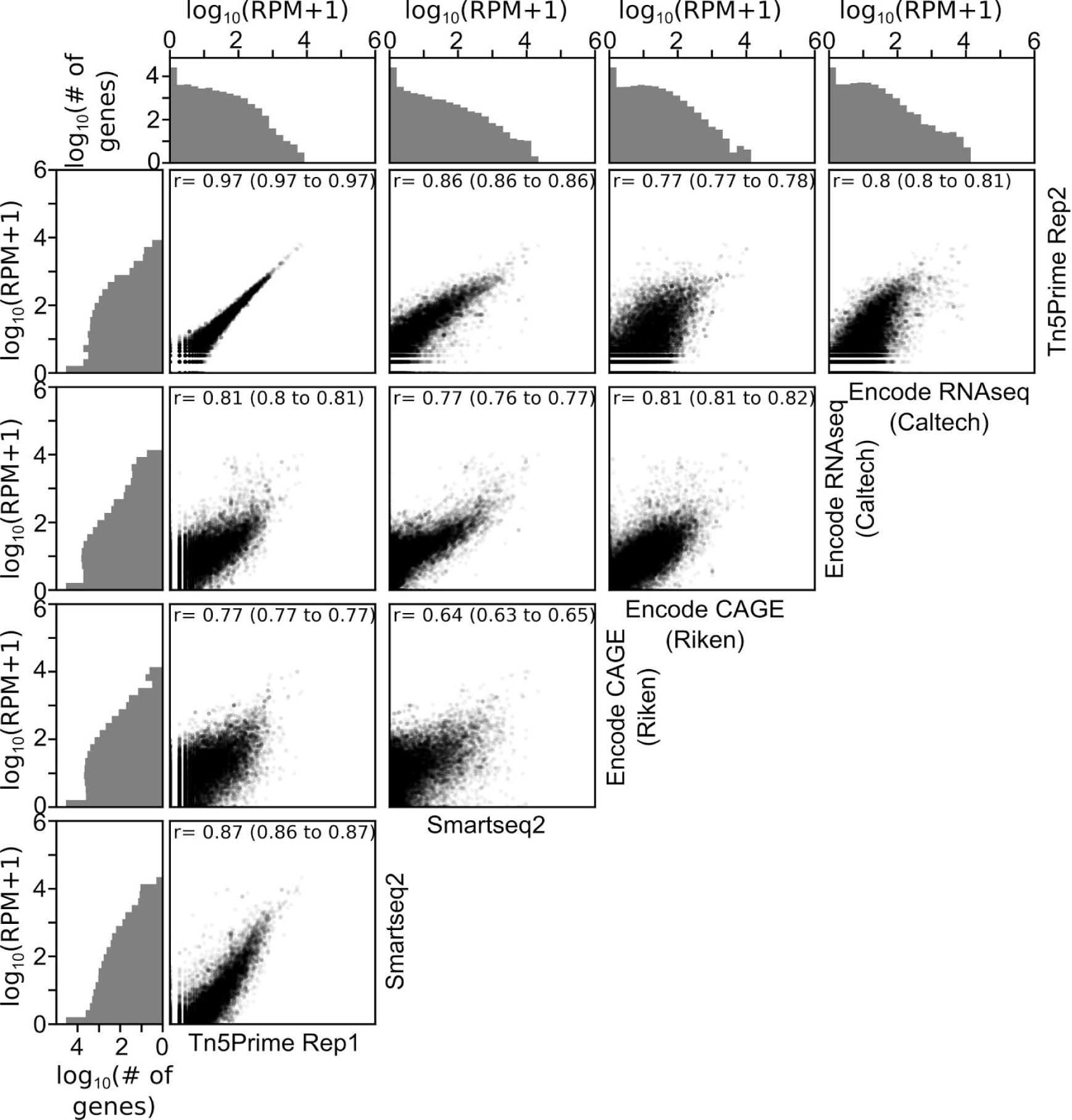
Tn5Prime quantifies transcriptomes accurately and reproducibly. Pairwise correlations of transcript levels between Tn5Prime, Smartseq2, ENCODE CAGE and ENCODE RNAseq are shown as scatter plots. Each transcript is shown as a black dot with an opacity of 5%. Distribution of transcript levels is shown on the outside of the plots in grey histograms.

### Transcript quantification and transcription start site localization in single B cells

As the Tn5Prime protocol is based on the same cDNA amplification strategy as the Smartseq2 protocol, we expected it capable of generating sequencing libraries from single cells. Indeed, we successfully generated single cell libraries using the Tn5Prime protocol from primary murine B-lymphocytes (B2 cells; IgM+B220+CD5-CD11b-)(n=12) isolated from the peritoneal cavity. We generated between 17,534-93,429 2x300 bp read pairs per cell using the Illumina MiSeq of which 62% passed quality filtering. Of the filtered reads, an average of 91.48% uniquely mapped to the mouse genome. The high alignment percentage indicates we are able to generate high quality libraries from single cells using our Tn5Prime. Despite the very low total number of read pairs we collected, we still detected 339 expressed genes per cell on average. These results are not surprising as primary B cells can contain little RNA [ref?] and transcript abundance in single cells can be substantially variable depending on the state of the cell [ref?]. Among the genes expressed in many of the single cells were genes corresponding to B cell function, including CD19, CD79a and components of the MHC complexes (Fig. S1) These data indicate that we can efficiently identify cell type specific genes.

### Analysis of 192 Single CD27^high^ CD38^high^ Human B Cells

After successfully testing our Tn5Prime method on single mouse B cells, we next wanted to develop a multiplex approach capable of evaluating hundreds of human single cells. To this end, we FACS sorted into 192 single B cells into individual wells of 96 well plates using the canonical surface molecules CD19, CD27 and CD38 to sub-select the plasmablast subpopulation (Fig. S2). Plasmablasts are one of the most widely studied B cell populations and are frequently monitored after vaccination or infections by flow cytometry. The plasmablast cell compartment is defined by high levels of surface markers CD27 and CD38, but separation from memory B cells which also express these markers, albeit at lower levels, can be challenging. Therefore, analyzing these cell types at the single cell level should help further delineate these populations.

By inserting cellular indexes into the template switch oligo during reverse transcription to pool libraries after reverse transcription. This allowed us to streamline our method and increase our throughput by decreasing the PCR and Tn5 reactions required. Using our multiplexing strategy we generated Tn5 libraries for 192 single B cells using 192 RT reactions, 24 PCR reactions and 24 Tn5 reactions. Although this was not performed, library pools carrying distinct Illumina sample indexes could have been further pooled following PCR to reduce the numbers of Tn5 reactions to 2.

We generated 194,553,648 150 bp paired end reads total. To determine gene expression for each cell, reads were assigned to one of 192 single cells based on its Illumina index reads and by comparing the sequence of the first 8 bases of read 1 to the cellular index sequences. 91% of the 194,553,648 150bp paired end reads were successfully assigned to one of the 192 single B cells. 90.75% of cell-assigned reads were successfully aligned to the human genome using STAR with a median of 74.59% percent of cell-assigned reads being uniquely assigned to an annotated gene. Each cell expressed a median of 534 genes. Of the 58234 annotated genes in GENCODE, 5414 genes had at least one read per cell on average. The median redundancy for each cell is 13.92 which means that, on average, each uniquely aligned cDNA fragment was sequenced 13.92 times. This indicates that the libraries were sequenced exhaustively.

### Detecting Transcription Start Sites in single CD27^high^ CD38^high^ B cells using Tn5Prime

To determine if transcription start site specificity is maintained within the single cell data, read 1 start distribution was compared to annotated transcription start sites and Encode CAGE data. By calling peaks, we found that our single cell results were able to maintain transcription start site specificity, with peaks predominantly falling within the annotated transcription start sites (Fig. 4A-B). In addition to the transcription start site, the directionality of transcription can be inferred due to our custom template switch oligo incorporating a forward-read priming site to the 5’ region of the transcript which is an advantage over many other single cell RNAseq protocol (Fig. 4C,D).

**Fig 4.**
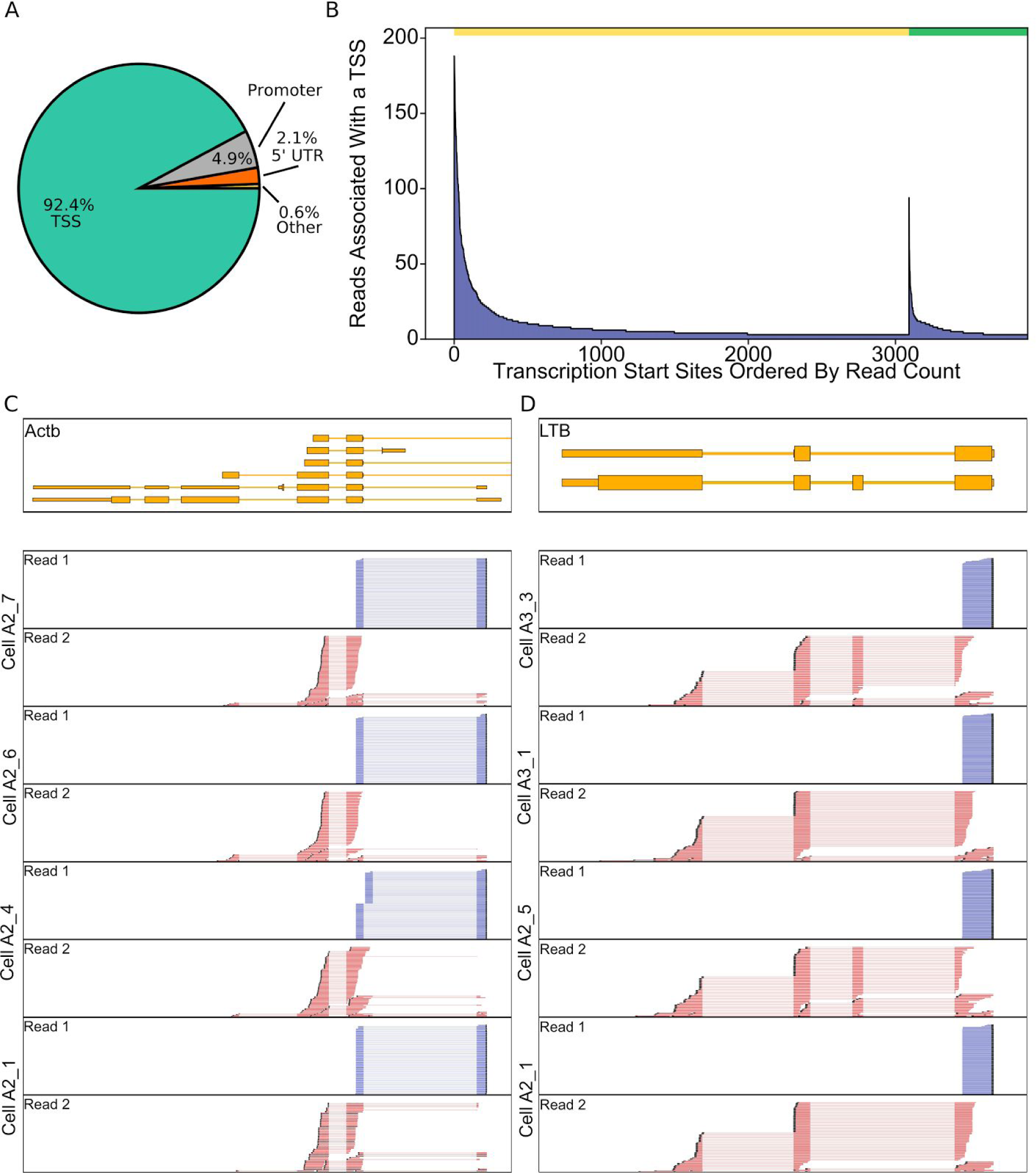
Transcription start sites are detected in single CD27^high^ CD38^high^ B cells. A) CD27^high^ CD38^high^ Tn5Prime peaks were matched to features in the Gencode annotation and the feature they matched are shown as a pie chart. TSS = on or less than 25bp behind the start of an annotated GENCODE gene, 5’UTR = inside 5’ prime untranslated region, Promoter = between 26 and 1000bp behind start of annotated gene. B) Tn5 peaks were categorized into two groups. One group contains all peaks within 25bp of a peak identified in the complete RIKEN CAGE peak Human peak database and the other group contains all other peaks. These peaks were sorted by the number of cells associated with that peak in the CD27^high^ CD28^high^ B cell data set and displayed in figure 5a. The yellow bar indicates the peaks within 25bp and the green bar indicates all other peaks. C,D) Genome Browser view of reads of several cells aligned to Actb (C) and LTB (D) genes. In addition to TSS information, read alignments also show differential isoform usage between cells.

### Detecting Subpopulations within CD27^high^ CD38^high^ B cells using Tn5Prime

Since separating memory B cells and plasmablasts by FACS based on surface markers can be challenging, especially when the adaptive immune system is not perturbed, we wanted to see whether we could do so post-sorting using their gene expression profiles. Cells outside more than three median absolute deviations from the median for percent alignment, Mitochondrial transcript percentage, or number of detected genes were marked as outliers and eliminated prior to normalization of transcript counts (Fig. S3). After normalizing raw gene expression counts and removing non-recombined and therefore non-applicable antibody gene segments from the annotation (10), we clustered the remaining 159 sorted B cells using t-SNE dimensional reduction. The clusters were robust when the data was subsampled to 100,000 reads per cell (Fig. S4). We then identified genes that showed significant differences between the two clusters. We detected 411 genes with significant changes including J-chain, LTB, XBP-1, HSPA5, MZB1, as well as HLA-DRA, HLA-DRB5, and HLA-DPB1 (Table S2). J-chain was upregulated in cluster 2 and is involved in antibody secretion of IgM and IgA (11) (Fig. 5). We also found XBP-1, MZB1 and HSPA5 were upregulated within cluster 2 and are known targets of BLIMP-1 which is essential in plasmablast differentiation (Fig. S5) (12). LTB was downregulated in cluster 2 and has been shown to be downregulated upon B cell activation (13) (Fig. 5). HLA-DRA, HLA-DRB5, and HLA-DPB1 which encode for the alpha and beta chains of the MHC II complex were also downregulated in cluster 2, indicating less MHC II presentation to T cells which is indicative of plasma cells and plasmablasts (14)). Together, this suggests that cluster 2 does represent activated plasmablasts which are known to secrete more antibody and display less MHC II complex than the memory B cells in cluster 1.

**Figure 5.**
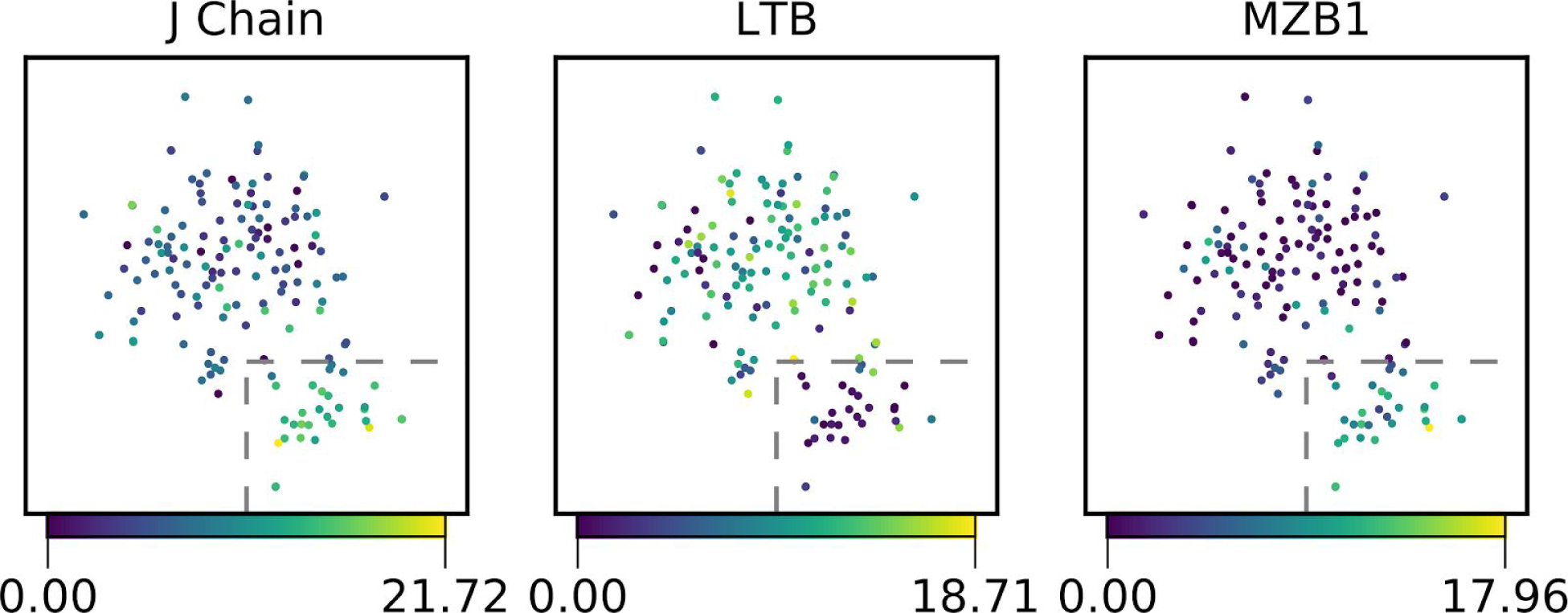
Clustering of CD27^high^ CD38^high^ B cells. 159 B cells were divided into two populations by t-SNE dimensionality reduction (15). In the three subplots, cells are colored based on their expression of example genes that were significantly differentially expressed between the two populations as determined by a multiple hypothesis testing corrected Mann-Whitney U tests. The cells inside the boxed area belong to cluster 2 and all other cells belong to cluster 1.

### Assembly of antibody heavy and light chain sequences from single B cell Tn5Prime data

Ideally, we would not only want to identify plasmablasts based on their gene expression profile, but also determine the sequences of the antibodies they express. Sequencing antibodies has been a long-standing challenge in B cell biology and antibody engineering because it requires the identification of unique pairs of rearranged antibody heavy and light chains for each cell. Current techniques rely either on the targeted amplification and sequencing of antibody heavy and light chain genes (16) in single cells or on the assembly of their sequences from non-targeted RNA-seq data (17). In contrast to 3’ end based Drop-Seq and 10X Genomics data, 5’ based Tn5Prime could potentially provide this antibody sequence information in addition to genome wide expression profiling, because the 5’ region contains the unique V(D)J rearrangement of heavy and light chain transcripts.

To determine if our Tn5Prime protocol could be used for assembling antibody heavy and light chain sequences, we assembled whole transcriptomes using SPAdes (18). IgBLAST (19) was used to identify transcripts containing V, D, and J gene segments rearranged in a productive manner. These transcripts were aligned on to Constant gene segments to identify isotype. The list of putative antibodies was then filtered for obvious cross-contamination and mis-assemblies. In this way, we effectively determined heavy and light chain sequences and identify their unique pairings within single B cells (Fig. 6A).

**Figure. 6.**
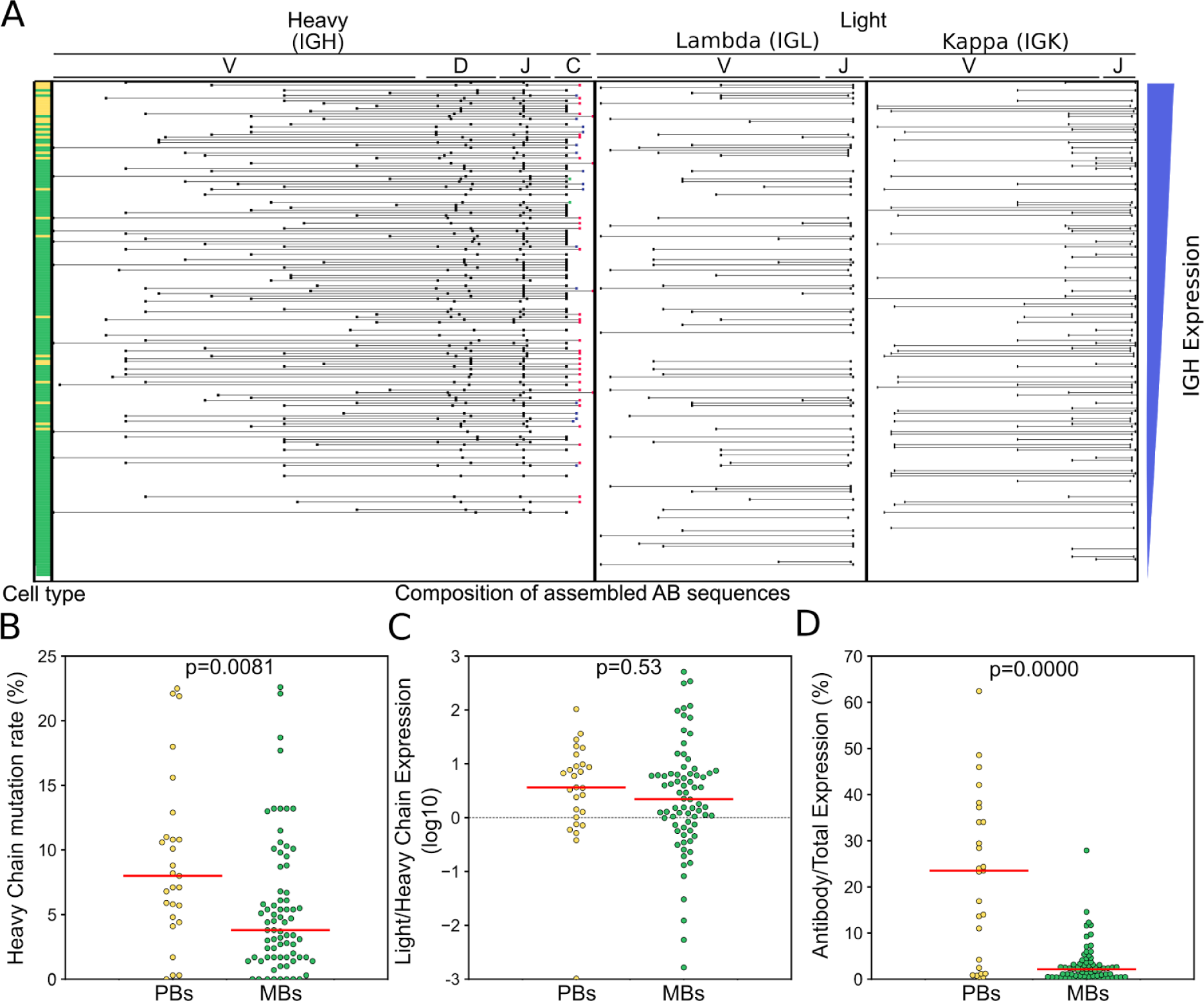
Assembling Antibody transcripts from Tn5Prime data. Antibody transcripts were assembled by generating complete assembled transcriptomes for each cell with SPADES and then using IGBLAST to search for transcripts with antibody features. Antibody transcripts for each cell were filtered for mis-assemblies and mis-annotations. Cells were sorted by the abundance of heavy chain transcripts in their Tn5Prime data and V(,D,) and J segment information for their heavy and light chains are shown in the schematic in the center. The putative cell type determined by clustering with t-SNE is indicated on the left. Yellow: plasmablasts, Green: Memory B cells. B-D) Antibody usage and characteristics were compared between plasmablasts and memory B cells. Somatic Hypermutation rates (B), light to heavy chain expression ratios (C) and the percentage of all aligned sequencing reads that originated from antibody transcripts (D) were compared using dotplots. Yellow: plasmablasts, Green: Memory B cells. Medians are shown as red lines. All p-values are calculated using two-sided Monte Carlo permutation test with 10000 permutations.

Of the 192 B-cells we analyzed, we were able to assemble one heavy chain and one light chain to 117 B-cells. Of these 117 B-cells 46 cells had a Lambda light chain and 71 cells had a Kappa light chain. Five additional cells had one heavy chain and two light chains, 35 cells had no heavy chains but at least one light chain, and 35 cells had no heavy chains and no light chains. To determine the sequencing depth requirement for successful heavy and light chain assembly, subsampling was performed on the reads and the assembly and pairing analysis redone (Fig. S6). We found 100,000 reads per cell was sufficient to assemble one heavy and one light chains for 91 of 117 B cells with successfully assembled chain pairs without subsampling.

101 and of the 117 cells with paired heavy and light chains also passed all other quality filters and were clustered by t-SNE into the putative plasmablast and memory B cell clusters. This combination of single cell identity and paired antibody sequences allowed us to perform detailed analysis of differences in antibody usage and characteristics between those two populations. First, the putative plasmablast population featured less IgM antibodies than the memory B cell population (19% IgM in plasmablasts vs 53% in memory B cells). Second, using IgBlast (19), we found that both heavy (Fig. 6B) and light chain sequences showed significantly higher levels of somatic hypermutation in plasmablasts than memory B cells (Heavy chain: median 8.0% vs 3.8% somatic hypermutation, two-sided Monte Carlo permutation test p-value=0.0081; Light chain: median 4.9% vs 2.7% somatic hypermutation, two-sided Monte Carlo permutation test p-value=0.0117). Third, by counting and normalizing sequencing reads originating from antibody transcripts, we could determine and compare heavy and light chain expression in these two populations. Generally, light chains were expressed about 3-fold higher than heavy chains (Fig. 6C) with no significant difference between plasmablasts and memory B cells (two-sided Monte Carlo permutation test p-value=0.533). However, the percentage of all aligned sequencing reads that originated from antibody transcripts showed dramatic differences between plasmablasts and memory B cells. The median percentage of reads that originated from antibody transcripts was 23.5% in plasmablasts and only 2.2% in memory B cells (Fig. 6D) (Monte Carlo Permutation test two-sided p-value=0). In one plasmablast over 60% of all aligned sequencing reads originated from antibody transcripts indicating just how much of the plasmablast transcriptome can be dedicated to the production and secretion of antibodies. In summary, our analysis of antibody usage and characteristics showed that plasmablasts express more mutated and class-switched antibodies at much higher levels than memory B cells.

## Discussion

Here we present a novel method for the genome-wide identification of transcription start sites in bulk samples and single cells. The method combines aspects of Smartseq2 and STRT. By modifying template-switch oligos used during reverse transcription to carry one sequencing adapter and loading the other sequencing adapter on the Tn5 enzyme used for cDNA fragmentation we anchor the sequence priming sites for read 1 of an Illumina read pair to the 5’ end of transcripts without the need for fragmentation, ligation, and enrichment steps. The resulting workflow is easy to implement and capable of generating hundreds of libraries within a day. An important feature of our Tn5Prime method is the option to integrate cellular indexes during reverse transcription and Illumina sample indexes during PCR before Tn5 tagmentation. This allows the pooling of samples early in the workflow and thereby reduces experiment complexity and reagent costs.

We validated the Tn5Prime protocol on both bulk RNA and single cells. First, using 5ng of total RNA from the GM12878 cell line, we yielded similar results as the ENCODE CAGE data with respect to the identification of transcripts start sites. However, the CAGE protocol used by the ENCODE consortium used several order of magnitude more RNA. As the Smartseq2 protocol is already widely used, we expect that the Tn5Prime assay with its similar workflow and low RNA input has the potential to become a valuable tool for transcriptome annotation and quantification in the RNA-seq toolbox.

In addition to the analysis of bulk samples, we show that our Tn5Prime method can be utilized for interrogating single cells, both human and mouse. The TSO-based multiplexing approach we implemented makes it possible to efficiently analyze thousands of cells, thereby increasing the throughput of plate based RNAseq library protocols in a manner that is straightforward and economical.

In contrast to other droplet or microwell based protocols, which interrogate only the 3’ ends of transcripts, the Tn5Prime protocols interrogates the 5’ ends of transcripts, thereby capturing the unique sequence information of adaptive immune system receptors expressed on B and T cells. These receptors are often hard to assemble due to their unique genomic rearrangement. Our data shows that by limiting sequencing reads to the 5’ end of transcripts we can analyze both transcriptomes as well as paired antibody heavy and light sequences with the low sequencing coverage of ~100,000 reads per cell, thereby enabling the analysis of thousands of B cells in a single sequencing run. This approach should, without any modification, also be applicable to T cells to map rearrangement of the T cell receptors. This can provide novel insights into the composition of B and T cell malignancies as well as the state and composition of the adaptive immune system with regards to solid tumors.

To highlight the power of our approach we isolated 192 single human B cells from PBMCs using canonical plasmablast markers. Not only were we able to assemble paired antibody transcripts, but we succeeded in clustering the cells into two populations based on their gene expression profiles. The genes differentially expressed between those clustered suggested their putative cell types. Cells in the putative plasmablast cluster expressed more XBP-1 (X-box binding protein 1), J-chain, HSPA5, and MZB1 (Marginal Zone B1), which are all involved in either B cell activation or antibody production and secretion. Consistent with less antigen presentation, cells in the putative plasmablast cluster also expressed less MHC II transcripts including HLA-DRA, HLA-DRB5, and HLA-DPB1. Finally, MS4A1 (CD20) is also expressed less in the cells of the putative plasmablast cluster and is known to be downregulated in activated B cells. Overall, this clearly established that we could distinguish activated, antibody secreting plasmablasts from resting, antigen presenting memory B-cells; cell-types which are difficult to distinguish using conventional FACS analysis.

In addition to cell-types, we showed that Tn5Prime can be used to determine individual B cells’ paired antibody sequences. Together, these data allowed us to compare antibody usage in plasmablasts and memory B cells, showing that plasmablast expressed higher levels of more mutated and class-switched antibodies. In addition to providing functional insight into cell populations, this information will make it possible to make informed decisions as to which antibody sequences could be further cloned and tested functionally for clinical, diagnostic, and research applications.

In summary, Tn5Prime is an RNAseq library construction protocol with a streamlined workflow that surpasses the economy and throughput of other plate-based protocols. While not reaching the throughput of droplet- and microwell-based protocols, it generates high quality data that enables the identification of transcription start sites and could be useful for analyzing 5’ UTR features or help improve incomplete genome annotations. Finally, Tn5Prime presents the currently highest throughput mechanism to comprehensively analyze the individual cells of the adaptive immune system by determining both paired adaptive immune receptor sequences and gene expression profiles.

## Methods

### Cell purification, RNA isolation and sorting

#### GM12878

RNA from 500,000 GM12878 cells was extracted using the RNeasy kit (Qiagen) according to manufacturer’s instructions.

#### Murine B2 cells

Mice were maintained in the UCSC vivarium according to IACUC-approved protocols. Single murine Ter119-CD3-CD4-CD8-B220^+^ IgM^+^CD11b^−^ CD5^−^ B2 cells were isolated from wild-type C57Bl/6 mice by peritoneal lavage and incubated with fluorescently-labeled antibodies prior to sorting. The following antibodies were used to stain B-cells: Ter119, CD3 (Biolegend; 145-2C11), CD4 (Biolegend; GK1.5), CD8a (Biolegend; 53-6.7), B220 (Biolegend; RA3-6B2), IgM (Biolegend; RMM-1), CD5 (Biolegend; 53-7.3), and CD11b (Biolegend; M1/70). Cells were analyzed and sorted using a FACS Aria II (BD), as described (20–22).

#### Human B cells

Primary human cells were collected from the blood of a fully consented healthy adult in a study approved by the Institutional Review Board (IRB) at UCSC. Single human B cells were isolated from PBMC using negative selection using RosetteSep (StemCell). The resulting B cells were sorted for CD19^+^ CD27^high^ and CD38^high^. The following antibodies were used for staining B cells: CD19 (BD Pharmingen; HIB19), CD27 (Biolegend; 0323), and CD38 (Biolegend; HB-7). Cells were sorted using FACS Aria II (BD) and analyzed using FlowJo v10.2 (FlowJo, TreeStar Software, Ashland, OR).

Both murine and human single cells were sorted into 96 well plates and directly placed into 4ul of Lysis Buffer - 0.1% Triton X-100, 0.2ul of SuperaseIn (Thermo), 1ul of oligodT primer (IDT), 1ul of dNTP (10mM each)(NEB) - and frozen at −80°C.

### RNA-seq library construction and sequencing

4ul of RNA or Single Cell Lysate was reverse transcribed using Smartscribe Reverse Transcriptase (Clontech) in a 10ul reaction including either a Smartseq2 TSO (Smartseq2 libraries) or a Nextera A TSO (Tn5Prime libraries) according to manufacturer’s instructions at 42°C. The resulting cDNA was treated with 1 ul of 1:10 dilutions of RNAse A (Thermo) and Lambda Exonuclease (NEB) for 30min at 37°C. The treated cDNA was amplified with KAPA Hifi Readymix 2x (KAPA) using the ISPCR primer and a Nextera A Index primer (Tn5Prime only).

The resulting PCR product was treated with Tn5 enzyme (23) loaded with either Tn5ME-A/R and Tn5ME-B/R (Smartseq2) or Tn5ME-B/R adapters only (Tn5Prime).

The Tn5 treated PCR product was then size selected using a E-gel 2% EX (Thermo) to a size range of 400-1000bp. GM12878 RNA Smartseq2 and Tn5Prime libraries were sequenced on an Illumina HiSeq2500 2x150 run, mouse B2 cell Tn5Prime libraries were sequenced on a Illumina MiSeq 2x300 run, and human B cell Tn5Prime libraries were sequenced on two Illumina HiSeq3000 runs.

### Sequencing alignment and analysis

Smartseq2, Tn5Prime, ENCODE CAGE (GEO accession GSM849368; produced by the lab of Piero Carnici at RIKEN), and ENCODE RNAseq (GEO accession GSM958742; produced by the lab of Barbara Wold at Caltech) (24) GM12878 data as well as Tn5Prime B2 data were trimmed of adapters low quality bases using trimmomatic (v0.33) (8) and a quality cutoff of Q15. Trimming of the 192 human B cell data was performed by Cutadapt, filtering out all paired reads where one or more read had a post-trimming length of less than 25 bp.

Trimmed reads derived from the GM cell line and single B cells were aligned to the human genome (GRCh38) annotated with Ensembl GRCh38.78 GTF release using STAR (v2.4) (9). Trimmed reads derived from the B2 cells were aligned to the mouse genome (GRCm38) annotated with Ensembl GRCm38.80 GTF release using STAR (v2.4). Expression levels were quantified using featureCounts (v1.4.6-p1) (25) and normalized by total read number resulting in RPM (<u>R</u>eads <u>P</u>er <u>M</u>illion).

Peaks for CAGE and Tn5Prime data were called by counting the number of unique fragments which began their forward read alignments (R1 for Tn5Prime) at each position within each chromosome and for each orientation (positive or negative). A peak was called at a position and orientation if at least five alignments begin at that position, the position one nucleotide downstream has fewer alignments beginning at that position, and the position one nucleotide upstream has fewer alignments beginning at that position. For the single cell data, peaks were filtered out unless they appeared in more than one cell. The distance between the TN5 peaks and the nearest CAGE peak was called by inserting the nucleotide coordinates of the CAGE peaks into kd-trees and then performing a nearest neighbor search on them using the TN5 peak coordinates. Each chromosome and orientation had its own kd-tree.

### Antibody Assembly

After assignment, reads were assembled into transcriptomes using rnaSPAdes (18) with the single-cell parameters. Putative immunoglobulin transcripts are detected and annotated by running IGBLAST (19) against the assembled transcriptome using Human V,D and J segments from the IMGT database (26). Isotypes are assigned to putative IG transcripts by aligning constant regions to the transcripts with BWA-MEM (27).

Antibody transcripts were filtered with the following process:

1. A table is generated from the SPADES/IGBLAST/BWA pipeline listing each putative IG transcript for each cell in which each row represents one assembled antibody transcript and contains information indicating which cell it came from, the overall abundance(as determined by BWA) within the cell,the CDR3 sequence and the type(IGH,IGK,IGL) as well as the inferred segments used during VDJ recombination.
2. The transcripts are clustered by CDR3 sequencing similarity using a single-linkage clustering algorithm Based on the Levenshtein distance where two clusters of transcripts are merged when at least one transcript CDR3 has a Levenshtein distance of 2 or less with the CDR3 of any transcript in another cluster.
3. Transcripts belonging to the same cluster are merged so that highly similar transcripts belonging to the same cell are combined and their counts added together. This is done to correct for the production of spurious alternative assemblies produced by SPADES within each cell's assembled transcriptome.
4. a list is generated for each transcript of the cells in which they appear.
5. The lists is sorted by the abundance of the transcript within the cells.
6. the entries in the lists are marked by their relative abundance. If the number of reads aligned to the transcript in a cell is less than 10% of the largest amount reads aligned to that transcript within any cell, it is marked as being a potential contaminant. The idea is that if a transcript discovered in a cell is a contaminant it should have at least an order of magnitude fewer reads associated with it when compared with the cell it actually came from.
7. For each type (IGH,IGK,IGL) of IG transcript found within each cell, the largest unique (non-contaminant) transcript is picked to have potentially come from that cell. if a unique transcript cannot be found, the most highly expressed transcript is selected
8. If both a potential IGK and IGL are present within a cell, the unique transcript is selected. if both are unique or non-unique the most highly expressed transcript is selected unless either transcript has an abundance of at least 10% of the other.
9. After this process, most cells should have a single heavy chain and a single light chain.

### Visualization

All data visualization was done using Python/Numpy/Scipy/Matplotlib (28–31). Schematics were drawn in Inkscape (https://inkscape.org/en/).

### Data and Script Access

Raw data has been uploaded to the Sequence Read Archive (SRA) under the accessions PRJNA320873 (GM12878 Smartseq2 and Tn5Prime), PRJNA320902 (Mouse B2 Cells), and PRJNA415475 (Human CD27^high^ CD38^high^). A UCSC genome browser track is available at

https://genome.ucsc.edu/cgi-bin/hgTracks?hgS_doOtherUser=submit&hgS_otherUserName=chkcole&hgS_otherUserSessionName=TN5_Prime_Alignments

The Tn5Prime and CAGE Peak Caller and peak distance calculator are available at https://github.com/chkcole/Peak-Calling. All other Scripts are available upon request.

## Acknowledgements

We thank the Sanford lab at UCSC for providing GM12878 cells and the Dubois lab at UCSC for expert help with producing Tn5 enzyme. This work was supported by an NIH/NIDDK award (R01DK100917) and an Alex’s Lemonade Stand Foundation Innovation Award to ECF; by CIRM Training grant TG2-01157 to AEB; and by CIRM Shared Stem Cell Facilities (CL1-00506) and CIRM Major Facilities (FA1-00617-1) awards to UCSC. ECF is the recipient of a California Institute for Regenerative Medicine (CIRM) New Faculty Award (RN1-00540) and an American Cancer Society Research Scholar Award (RSG-13-193-01-DDC). CV is a recipient of the 2017 Hellman Fellowship. A.B. and C.C. are funded by the NHGRI/NIH training grant 1T32HG008345-01.

## References

1. Picelli,S., Faridani,O.R., Björklund,A.K., Winberg,G., Sagasser,S. and Sandberg,R. (2014) Full-length RNA-seq from single cells using Smart-seq2. Nat. Protoc., 9, 171–181.

2. Treutlein,B., Brownfield,D.G., Wu,A.R., Neff,N.F., Mantalas,G.L., Espinoza,F.H., Desai,T.J., Krasnow,M.A. and Quake,S.R. (2014) Reconstructing lineage hierarchies of the distal lung epithelium using single-cell RNA-seq. Nature, 509, 371–375.

3. Darmanis,S., Sloan,S.A., Zhang,Y., Enge,M., Caneda,C., Shuer,L.M., Hayden Gephart,M.G., Barres,B.A. and Quake,S.R. (2015) A survey of human brain transcriptome diversity at the single cell level. Proc. Natl. Acad. Sci. U. S. A., 112, 7285–7290.

4. Islam,S., Kjällquist,U., Moliner,A., Zajac,P., Fan,J.-B., Lönnerberg,P. and Linnarsson,S. (2011) Characterization of the single-cell transcriptional landscape by highly multiplex RNA-seq. Genome Res., 21, 1160–1167.

5. Islam,S., Zeisel,A., Joost,S., La Manno,G., Zajac,P., Kasper,M., Lönnerberg,P. and Linnarsson,S. (2014) Quantitative single-cell RNA-seq with unique molecular identifiers. Nat. Methods, 11, 163–166.

6. Salimullah,M., Sakai,M., Mizuho,S., Plessy,C. and Carninci,P. (2011) NanoCAGE: a high-resolution technique to discover and interrogate cell transcriptomes. Cold Spring Harb. Protoc., 2011, db.prot5559.

7. Shiraki,T., Kondo,S., Katayama,S., Waki,K., Kasukawa,T., Kawaji,H., Kodzius,R., Watahiki,A., Nakamura,M., Arakawa,T., et al. (2003) Cap analysis gene expression for high-throughput analysis of transcriptional starting point and identification of promoter usage. Proc. Natl. Acad. Sci. U. S. A., 100, 15776–15781.

8. Bolger,A.M., Lohse,M. and Usadel,B. (2014) Trimmomatic: a flexible trimmer for Illumina sequence data. Bioinformatics, 30, 2114–2120.

9. Dobin,A., Davis,C.A., Schlesinger,F., Drenkow,J., Zaleski,C., Jha,S., Batut,P., Chaisson,M. and Gingeras,T.R. (2013) STAR: ultrafast universal RNA-seq aligner. Bioinformatics, 29, 15–21.

10. Lun,A.T.L., Bach,K. and Marioni,J.C. (2016) Pooling across cells to normalize single-cell RNA sequencing data with many zero counts. Genome Biol., 17, 75.

11. Lamson,G. and Koshland,M.E. (1984) Changes in J chain and mu chain RNA expression as a function of B cell differentiation. J. Exp. Med., 160, 877–892.

12. Minnich,M., Tagoh,H., Bönelt,P., Axelsson,E., Fischer,M., Cebolla,B., Tarakhovsky,A., Nutt,S.L., Jaritz,M. and Busslinger,M. (2016) Multifunctional role of the transcription factor Blimp-1 in coordinating plasma cell differentiation. Nat. Immunol., 17, 331–343.

13. Zhu,X., Hart,R., Chang,M.S., Kim,J.-W., Lee,S.Y., Cao,Y.A., Mock,D., Ke,E., Saunders,B., Alexander,A., et al. (2004) Analysis of the major patterns of B cell gene expression changes in response to short-term stimulation with 33 single ligands. J. Immunol., 173, 7141–7149.

14. Calame,K.L., Lin,K.-I. and Tunyaplin,C. (2003) Regulatory mechanisms that determine the development and function of plasma cells. Annu. Rev. Immunol., 21, 205–230.

15. Maaten,L. van der and Hinton,G. (2008) Visualizing Data using t-SNE. J. Mach. Learn. Res., 9, 2579–2605.

16. Wrammert,J., Smith,K., Miller,J., Langley,W.A., Kokko,K., Larsen,C., Zheng,N.-Y., Mays,I., Garman,L., Helms,C., et al. (2008) Rapid cloning of high-affinity human monoclonal antibodies against influenza virus. Nature, 453, 667–671.

17. Canzar,S., Neu,K.E., Tang,Q., Wilson,P.C. and Khan,A.A. (2017) BASIC: BCR assembly from single cells. Bioinformatics, 33, 425–427.

18. Bankevich,A., Nurk,S., Antipov,D., Gurevich,A.A., Dvorkin,M., Kulikov,A.S., Lesin,V.M., Nikolenko,S.I., Pham,S., Prjibelski,A.D., et al. (2012) SPAdes: a new genome assembly algorithm and its applications to single-cell sequencing. J. Comput. Biol., 19, 455–477.

19. Ye,J., Ma,N., Madden,T.L. and Ostell,J.M. (2013) IgBLAST: an immunoglobulin variable domain sequence analysis tool. Nucleic Acids Res., 41, W34–40.

20. Ugarte,F., Sousae,R., Cinquin,B., Martin,E.W., Krietsch,J., Sanchez,G., Inman,M., Tsang,H., Warr,M., Passegué,E., et al. (2015) Progressive Chromatin Condensation and H3K9 Methylation Regulate the Differentiation of Embryonic and Hematopoietic Stem Cells. Stem Cell Reports, 5, 728–740.

21. Smith-Berdan,S., Nguyen,A., Hong,M.A. and Forsberg,E.C. (2015) ROBO4-mediated vascular integrity regulates the directionality of hematopoietic stem cell trafficking. Stem Cell Reports, 4, 255–268.

22. Beaudin,A.E., Boyer,S.W. and Forsberg,E.C. (2014) Flk2/Flt3 promotes both myeloid and lymphoid development by expanding non-self-renewing multipotent hematopoietic progenitor cells. Exp. Hematol., 42, 218–229.e4.

23. Picelli,S., Björklund,A.K., Reinius,B., Sagasser,S., Winberg,G. and Sandberg,R. (2014) Tn5 transposase and tagmentation procedures for massively scaled sequencing projects. Genome Res., 24, 2033–2040.

24. ENCODE Project Consortium (2012) An integrated encyclopedia of DNA elements in the human genome. Nature, 489, 57–74.

25. Liao,Y., Smyth,G.K. and Shi,W. (2014) featureCounts: an efficient general purpose program for assigning sequence reads to genomic features. Bioinformatics, 30, 923–930.

26. Lefranc,M.-P., Giudicelli,V., Ginestoux,C., Bosc,N., Folch,G., Guiraudou,D., Jabado-Michaloud,J., Magris,S., Scaviner,D., Thouvenin,V., et al. (2004) IMGT-ONTOLOGY for immunogenetics and immunoinformatics. In Silico Biol., 4, 17–29.

27. Li,H. (2013) Aligning sequence reads, clone sequences and assembly contigs with BWA-MEM. arXiv preprint arXiv:1303. 3997.

28. Hunter,J.D. (2007) Matplotlib: A 2D Graphics Environment. Comput. Sci. Eng., 9, 90–95.

29. Oliphant,T.E. (2007) Python for Scientific Computing. Comput. Sci. Eng., 9, 10–20.

30. van der Walt,S., Colbert,S.C. and Varoquaux,G. (2011) The NumPy Array: A Structure for Efficient Numerical Computation. Comput. Sci. Eng., 13, 22–30.

31. Jones,E., Oliphant,T. and Peterson,P. (2001--) {SciPy}: Open source scientific tools for {Python}.

